# Time-dependent fitness effects can drive bet-hedging populations extinct

**DOI:** 10.1101/054007

**Authors:** Eric Libby, William Ratcliff

**Affiliations:** Santa Fe Institute, Santa Fe, New Mexico, United States; School of Biology, Georgia Institute of Technology, Atlanta, Georgia, United States

**Keywords:** stochastic switching, time-dependent fitness, bet-hedging, competition, survival

## Abstract

To survive unpredictable environmental change, many organisms adopt bet-hedging strategies that trade short-term population growth for long-term fitness benefits. Because the benefits of bet-hedging may manifest over long time intervals, bet-hedging strategies may be out-competed by strategies maximizing short-term fitness. Here, we investigate the interplay between two drivers of selection, environmental fluctuations and competition for limited resources, on different bet-hedging strategies. We consider an environment with frequent disasters that switch between which phenotypes they affect in a temporally-correlated fashion. We determine how organisms that stochastically switch between phenotypes at different rates fare in both competition and survival. When disasters are correlated in time, the best strategy for competition is among the worst for survival. Since the time scales over which the two agents of selection act are significantly different, environmental fluctuations and resource competition act in opposition and lead populations to evolve diversification strategies that ultimately drive them extinct.

## Introduction

In the face of unpredictable environmental change, some organisms have evolved diversification strategies that generate offspring poorly suited to the current environment, but well-adapted to a different future environment [1, 2, 3]. For example, clonal bacterial lineages have evolved to produce both fast and slow growing phenotypes; the latter can better survive lethal antibiotic exposure [4, 5]. Organisms that adopt these strategies hedge their bets by sacrificing short-term increases in population expansion for potential long-term population growth [3, 6]. The time interval over which the long-term benefits of bet-hedging are realized depends in part on the frequency of environmental change [7, 8, 9]. Frequent environmental change can impose immediate selection against organisms who do not bet-hedge [10], while hedging against rare events may require long time periods [11]. Environmental change, however, will only rarely be the sole driver of selection. Other processes, such as competition for limited resources, may act on shorter time scales than environmental fluctuation, causing optimal bet-hedging strategies to go extinct before the long-term benefits are realized. In this paper, we investigate the interplay between these two drivers of selection.

Bet-hedging is a well-known survival strategy, evolved by diverse organisms, to increase fitness in risky, unpredictable environments [12, 13, 14, 15, 16, 17, 18, 19, 20]. For example, desert annuals delay germination in some seeds to hedge against across-year variation in spring rainfall [1, 21]. Another example is the bacterial pathogen *Haemophilus influenza*, in which a single clone generates offspring with diverse surface antigens that increase the probability that some of the population will avoid destruction by the host immune response [22, 23]. Concomitant with the abundance of bet-hedging, there are a wide array of molecular mechanisms for creating phenotypic diversity, including contingency loci, stochastic gene expression, developmental instability, and asymmetric cell division [24, 25, 26, 27]. These diversification mechanisms work together with reproductive strategies (e.g. asexual/sexual, clutch size) to enact particular forms of bet-hedging [12]. Considering the gamut of bet-hedging strategies is outside the scope of this paper, so for simplicity we will consider microbial bet-hedging, which is the focus of a large body of theoretical research [11, 28, 29, 30, 31, 32, 33].

Microbial bet-hedging is usually equated with stochastic switching strategies whereby a single genotype produces phenotypic heterogeneity in the absence of an apparent signal or regulatory response [10, 13, 24, 34, 35]. Mathematical models of microbial bet-hedging typically assume that the organism switches reversibly between at least two distinct phenotypic states, and that these distinct phenotypic states are each suited to different possible environmental states. Such models align well with experimental bet-hedging systems including CAP+/- phenotypes in *Pseudomonas fluorescens* [36], antigen expression in *Salmonella* [37, 38], competence to non-competence switch for DNA transformation in *Bacillus subtilis* [39], and galactose utilization in engineered populations of *Saccharomyces cerevisiae* [40]. For the purposes of modeling, phenotypic switch events are typically random and independent so that population-level heterogeneity follows a binomial or multinomial distribution, depending on the number of phenotypic states. A key area where models differ is the way in which environments fluctuate. For example, environmental fluctuations can occur randomly or after a fixed amount of time [28, 31, 32, 40, 41] and they can be symmetric or asymmetric in terms of how they switch and the selective pressure they exert [33, 41]. Whether or not stochastic switching confers a fitness benefit depends on the precise nature of environmental fluctuations.

In exponentially growing populations, the optimal rate of switching maximizes long-term geometric mean fitness [1, 8, 9, 42]. Indeed, models with expanding populations make it possible to calculate asymptotic growth rates [32]. The situation might be different if, along with environmental fluctuations, organisms face limitations in resources so that there exists a carrying capacity that restricts population size. If the population has not yet reached the carrying capacity, then each time an organism reproduces there is one less opportunity for reproduction by others in the population [43]. This couples the switching strategy of one organism to others, which could reward strategies that deny reproductive opportunities to other switching types either in the current environment or some future environmental state. Furthermore, the finite limit of population size allows for the possibility of extinction due to demographic stochasticity.

Here, we examine the consequences of coupling and extinction, introduced through a carrying capacity, on the evolution of microbial bet-hedging. We extend a previously published model [12] in which phenotypes experience periodic disasters such as might occur as the result of an adaptive immune response. We allow disasters to be correlated in time and determine how switching strategies fare in competition and survival, i.e. long-term fitness. When disaster risk is uncorrelated in time, the best competitive strategy also maximizes long-term survival. The situation changes when disaster risk is strongly correlated in time; the best strategy for competition is among the worst for survival. Our results show that environmental fluctuations and resource competition can lead to the evolution of diversification strategies that ultimately drive populations extinct.

## Methods

### Stochastic simulations

We consider a population of genotypes that switch between two phenotypic states, A and B. The defining characteristic of a genotype is its probability/rate of switching between phenotypes, which we denote as *p*. We assume that the switch occurs stochastically upon reproductive events so that each time an A or B reproduces there is a fixed probability *p* that it yields a cell of the opposite type. The phenotypic states do not differ in any fitness relevant trait other than susceptibility to risk. This risk manifests in disasters that target either A or B phenotypes and removes them completely from the population. We simulate the evolution of populations using a discrete time approach.

At each time step, there is a probability that a disaster occurs. We use a disaster probability of 10% for most results in this paper but explore the effects of changing this probability on time scales in Figure 5. If a disaster occurs, we then determine which phenotype it targets. A disaster targets the same phenotype as the previous disaster with probability *t*_*c*_ and the alternate phenotype with probability 1–*t*_*c*_ (the first disaster target is random). If *t*_*c*_ = .5 then the disaster targets a phenotype with no memory of previous targets. For *t*_*c*_ > .5 there is an increased chance that the phenotype targeted will be the same as before. In this way, the parameter *t*_*c*_ determines the duration of an environment hostile towards *A* or *B* phenotypes and the <u>t</u>emporal <u>c</u>orrelation of disasters.

Following the potential disaster, there is an opportunity for population turnover whereby a small fraction of the population is chosen for death. We use a default population turnover probability of 10% for most results in this paper but show the effects of changing this parameter in Figures 4 and 5. As a consequence of population turnover, even if a disaster does not occur the population still evolves.

After the effects of a possible disaster and population turnover, the remaining organisms reproduce until the population is restored to a fixed size, the carrying capacity *N*. For most results of the paper we consider *N* = 1000. Increasing this parameter increases the duration of competitions and survival. The effects of different values of *N* on Figure 4 are considered in the Supplementary material. Reproduction to carrying capacity occurs through an iterative process whereby organisms are randomly chosen to reproduce according to their frequency in the population. The discrete time step ends once the carrying capacity is reached. We simulate the populations until one genotype goes extinct or a maximum number of time steps occur, here 10^6^. Computer simulations were conducted in the programming language julia and are provided in the Supplementary material.

### Differential equation model

As a complement to our stochastic simulations we consider a deterministic model that uses differential equations (Eqns 1) to model population regrowth following a disaster. The simulation is still split into discrete rounds of disasters and regrowth however these events do not occur probabilistically. Instead, disasters occur every round and target one phenotype for a fixed number of times. For example, there may be five rounds of disasters targeting A types followed by five rounds of disasters targeting B types. After each disaster the population regrows for a fixed amount of time, (*t* = 100, though we consider other values of *t* in the Supplementary material), with a constant rate of population turnover described by the parameter *α*. We compute the frequency of types that switch according to probability *p*_1_ which is equal to 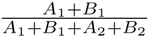 after 1000 environmental cycles. The differential equations were solved using Matlab and computer code is provided in the Supplementary material.

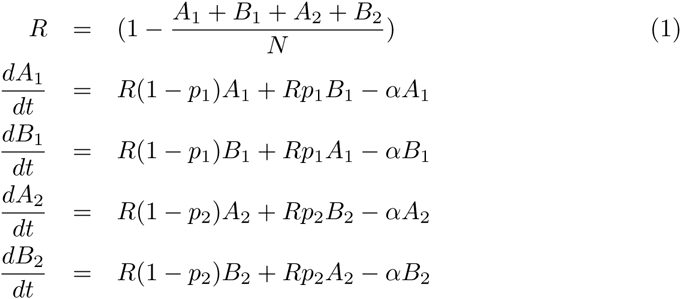

## Results

### Survival

In our model, the probability of switching phenotypes determines whether an organism can survive the challenge of repeat disasters. We vary the probability of repeat disasters (*t*_*c*_) from 0 to 1 and compute for each value of *t*_*c*_ the number of times out of 1000 stochastic simulations a switching strategy survives 10^6^ rounds of potential disasters, population turnover, and regrowth (see Figure 1). For fast switching probabilities, *p* ≥ .1, genotypes rarely go extinct (less than 1% of the time). In contrast, for slow switching probabilities, *p* ≤ .025, all organisms went extinct before the end of 10^6^ rounds. For switching probabilities in between these values, e.g. *p* = .05 and *p* = .075, there is a non-monotonic relationship between *t*_*c*_ and the frequency of extinction. Genotypes are less likely to go extinct at the extremes of *t*_*c*_, i.e. close to 0 or 1, as compared to intermediate values, e.g. *t*_*c*_ = .65.

**Figure 1:**
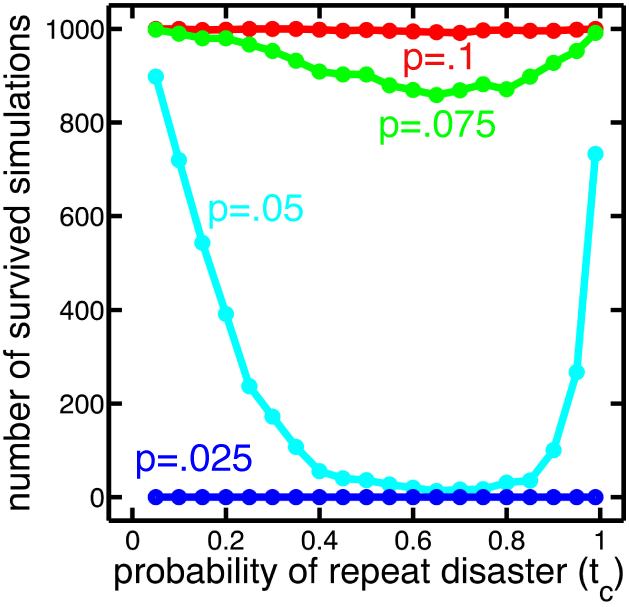
Survival as a function of the probability of switching and the probability of a repeat disaster. The number of extinctions out of 1000 for genotypes with different switch probabilities is shown as a function of the probability that a disaster repeats the phenotype it targets. Fast switchers *p* ≥ .1 rarely go extinct while slow switchers *p* ≤ .025 always go extinct. Intermediate values of p show a non-monotonic relationship in which extinction reaches a maximum around *t*_*c*_ = .65.

At *t*_*c*_ close to 1 there are long periods of disasters targeting the same phenotype. In the extreme case where *t*_*c*_ = 1 disasters never switch phenotypes, so as long as an organism diversified into two phenotypes at least once it would survive. As *t*_*c*_ decreases from 1 the probability of extinction increases because disasters more frequently switch between phenotypes.

At the opposite extreme, with *t*_*c*_ close to 0, disasters frequently switch between targeted phenotypes. While this seems like a challenging environment to survive, it is actually easier than an environment with longer periods of disasters targeting the same phenotype. The reason is that extinction in these simulations occurs through a lack of phenotypic diversification: an organism exists entirely in one phenotypic state when a disaster strikes that phenotype. After a disaster annihilates one phenotype, say *A*, the surviving phenotype, *B*, has a certain number of reproductive events in which to diversify. The probability that a *B* fails to produce at least one of the opposite phenotype, *A*, is (1 – *p*)^*m*^ where *p* is the switching probability and *m* is the number of reproductive events. The expected number of *A* types produced is *mp*, assuming that the *B* phenotype is the only one who gets to reproduce. Now, there are *N* – *mp* organisms of type *B* and *mp* of type *A*. This sets the stage for why an environment with *t*_*c*_ = 0 is easier to survive than one with an intermediate value of *t*_*c*_ = .65.

If the next disaster switches targets and annihilates the *B* type, as would be the case if *t*_*c*_ = 0, then the *A* types would have *N* – *mp* opportunities to diversify and produce a *B* type. The chance of failure here is (1 – *p*)^*N*–*mp*^. If, instead, *t*_*c*_ were greater than zero then there is a chance that the disaster would target the same phenotype as before, and annihilate any new *A* types produced. This would give the *B* phenotypes only *mp* opportunities to diversify and produce an *A*. The chance of failure in this case is (1 – *p*)^*mp*^. The difference between these two chances of failures can be many orders of magnitude: if, for instance, *p* = .05, *m* = 500, and *N* = 1000 then the probability of not diversifying goes up from 1.91 * 10^−20^% to 27.7%.

### Competition

If the only selective pressure were long-term survival, then the probability of switching should be high to avoid extinction. However, there are often other forms of selection acting on populations. We now consider what happens when there is competition in the form of another genotype present in the population. We vary the probability of repeat disasters (*t*_*c*_) from 0 to 1 and compute for each value of *t*_*c*_ which switching strategy is the best in pairwise competitions. The best strategy is the one that drives competitors extinct more often than it, itself, goes extinct. The optimal probability of switching decreases the more a disaster is likely to target the same phenotype, i.e. the higher the value of *t*_*c*_ (see Figure 2A). If the disasters frequently switch phenotypes such that *t*_*c*_ ≤ 0.50 then the best strategy in pairwise competitions is to rapidly diversify and switch phenotypes often. Thus, in this regime the optimal switching probability is *p* = 1.

**Figure 2:**
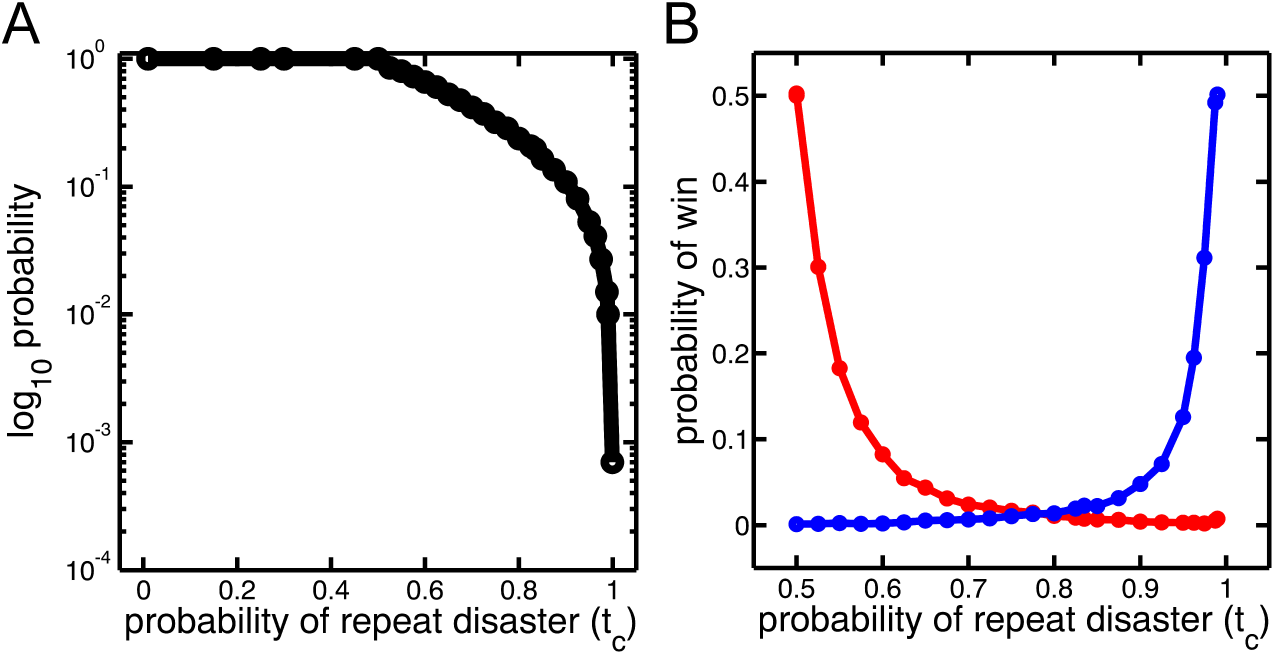
Optimal switching strategy versus the probability that a disaster targets the same phenotype. **A**) The switching probability that beats all others in pairwise competitions is shown as a function of the probability of repeat disasters *t*_*c*_. The switching probability decreases with increasing value of *t*_*c*_. **B**) A switch probability of *p* = 1 (red) or *p* = .01 (blue) is competed against the optimal switch probability for a range of *t*_*c*_ values (on the horizontal axis). Each strategy quickly drops in performance, as measured by the number of wins, by a factor of more than 5 with a .1 change in *t*_*c*_.

If, instead, disasters seldom switch the phenotype they target (*t*_*c*_ >> .5) then there is a cost to phenotypic diversification. Consider the case in which a disaster has removed all of the A phenotypes. As the B phenotypes reproduce to reach the carrying capacity any A types they produce will likely be lost to the next disaster. On the other hand, failing to diversify at all, *p* = 0, will lead to the genotype going extinct should the disaster switch the phenotype it targets. When risk is correlated in time, the optimal switching strategy must strike a balance between diversifying too much into the form that the disaster is targeting and not diversifying at all. As a point of reference from Figure 2A, if *t*_*c*_ = 0.99 then the optimal switch probability that strikes this balance is *p* = 0.01. These results echo earlier studies of bet-hedging populations in the absence of carrying capacity [3, 6, 44].

Although the optimal switching strategy changes with the probability of repeat disaster (*t*_*c*_), it is unclear how poorly a suboptimal switching strategy performs. To test this, we picked the best switching strategies for *t*_*c*_ = .5 (*p* = 1) and *t*_*c*_ = .99 (*p* = .01) and competed them against the optimal switching strategies for a range of *t*_*c*_ values (see Figure 2B). The performance quickly drops off such that if either strategy is competed against the optimal strategy at a *t*_*c*_ different by .1, it wins less than 10% of the time. Furthermore, *p* = 1 competes as unsuccessfully against *p* = .01 at *t*_*c*_ = .99 as *p* = .01 competes against *p* = 1 at *t*_*c*_ = .5 – they each win less than 1% of the time.

Due to the stochastic nature of these competitions, suboptimal strategies can occasionally beat optimal strategies. To understand what happens in these competitions, we investigate the competition between *p* = 1 and *p* = .01 at *t*_*c*_ = .99. On the rare occasions that *p* = 1 wins, the dynamics of disasters show frequent change in the targeted phenotype, mimicking an environment with a lower value of *t*_*c*_ in which *p* = 1 is more adaptive (see Figure 3A). In contrast, the more typical scenario is that disasters infrequently switch the target phenotype and thereby penalize strategies that adopt rapid phenotypic diversification (see Figure 3B). The trajectory of this extinction shows that each disaster gives an incremental numerical advantage to the slower switching strategy. This acts as a steady drain which ultimately leads the *p* = 1 genotype to extinction.

**Figure 3:**
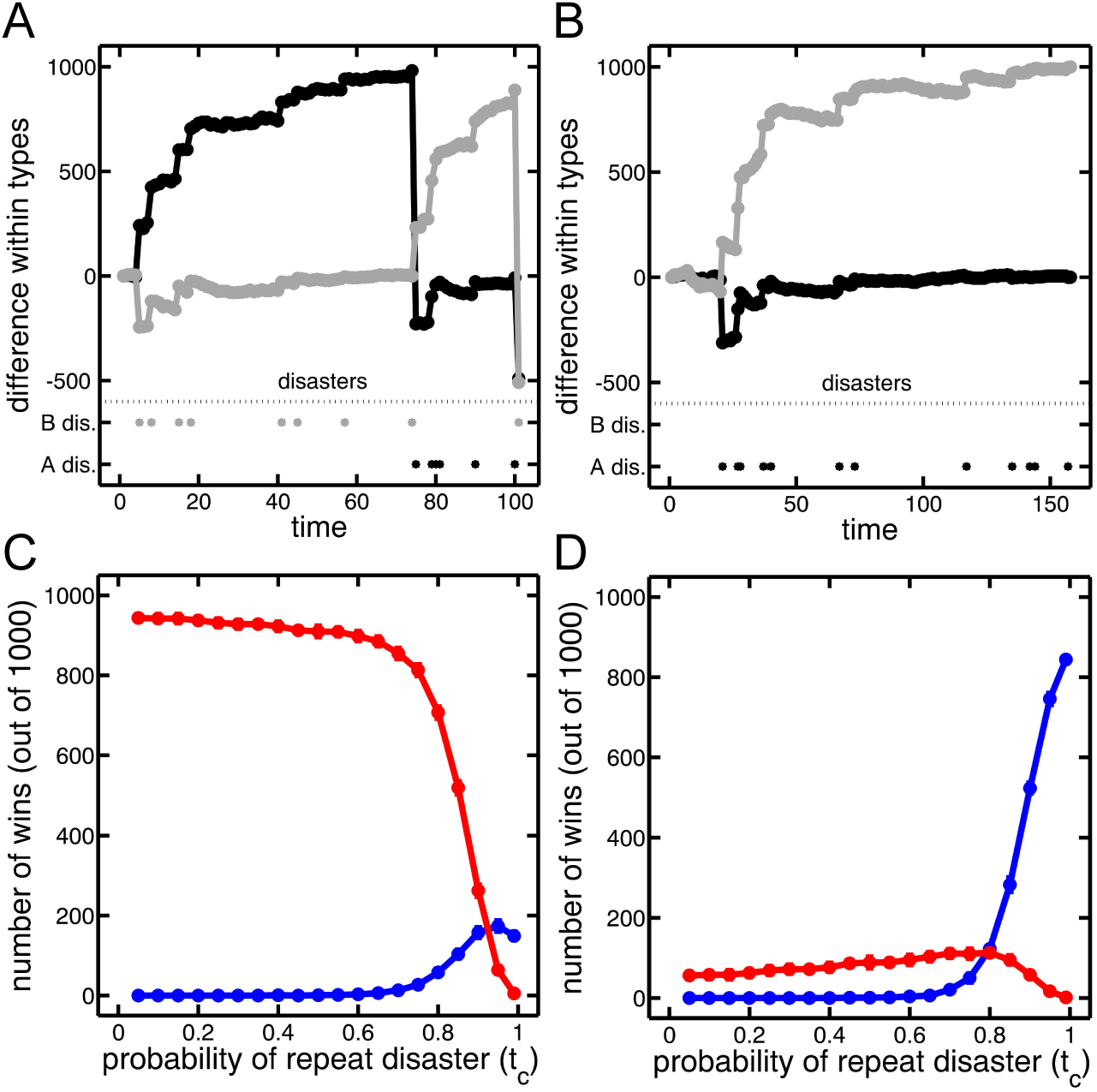
Characteristic manner different switching strategies win. **A**) The difference between A (gray) and B types (black) of a slow (*p* = .01) and fast (*p* = 1) switching strategy, e.g. *A*_slow_ - *A*_fast_, are plotted over the course of many disasters with *t*_*c*_ = .99. The phenotype targeted by the disaster is shown at the bottom. The faster switching phenotype wins because the disasters switch targets and mimic an environment with a lower *t*_*c*_. **B**) Here, the slow switching phenotype wins as repeated disasters slowly diminish the fast switching population. **C**) The number of wins (out of 1000) decided by a disaster switching phenotypes is shown as a function of the probability of repeat disasters. The fast switcher (*p* = 1, red) wins most of the competitions over the slow switcher (*p* = .01, blue) in this manner. **D**) Similar to C) except population turnover causes genotypes to win. In comparison to C), the slow switcher (*p* = .01, blue) wins more than 80% of its victories in this manner. Thus, fast switchers tend to win when disasters target both A and B phenotypes in rapid succession while slow switchers tend to win via a longer draining process.

The different trajectories in Figure 3A and 3B demonstrate the two ways in which organisms can go extinct during competition in our mathematical model. The first way is through a lack of phenotypic diversification as was discussed in the “Survival” section and is how the fast switcher beats the slow switcher (see Figure 3C). The second way is through population turnover and is how the slow switcher defeats the fast switcher (see Figure 3D). Extinction due to population turnover occurs during the death and replacement phase of our simulations. A consequence of replacement is that there are fluctuations in population abundances. If each organism has a probability *α* to be chosen for replacement, i.e. death, then the probability a population of *m* organisms goes extinct in a single round of replacement is *α*^*m*^. In order for this form of extinction to be realized a genotype must be rare, i.e. *m* must be small.

Although genotypes could become rare randomly through a set of unfortunate replacement events, usually it occurs because of a particular sequence of disasters. For instance, there could be a sequence of sudden switches in the phenotype targeted for disaster which would leave genotypes that switch infrequently as the rare type. Alternatively when disasters repeatedly target a single phenotype then the genotype that switches frequently can become rare. The latter case befalls the genotype with *p* = 1 in an environment with *t*_*c*_ = .99. To illustrate how this happens, consider two genotypes with switching probabilities *p*_1_ and *p*_2_ that just experienced a disaster eliminating all *m* of one phenotype. In the growth back to carrying capacity, we assume for simplicity that they evenly split the remaining spots in the population, i.e. 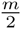 reproductive events are allotted to each genotype after the first disaster. Over *k* disasters and no replacement other than growth back to carrying capacity, then the genotype with a switch probability of *p*_1_ will gain an amount shown in Eqn. 2.

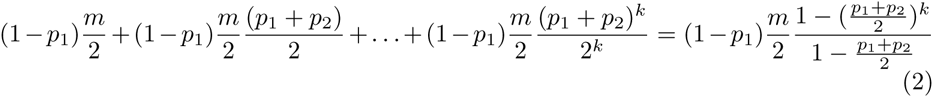

Thus, the genotype with *p*_1_ will have 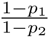 as much of the m pool as the genotype with *p*_2_. If one genotype switches with *p* = 1 then the other will eventually get the entire pool of *m*. This route to rarity is particularly effective if *m* is close to the carrying capacity *N*. Once a type is rare then population turnover can lead it to extinction.

The route to extinction that relies on population turnover is not unique to our stochastic simulation model. We can reformulate our model such that population growth occurs deterministically according to a set of differential equations (see Methods: Differential equation model, and Eqn. 1). These equations consider continuous population turnover rather than discrete rounds as was the case in the stochastic simulations. If populations grow for a fixed time according to these equations and then experience disasters that reset the A or B phenotypes back to 0 then we find that the amount of population turnover, i.e. the value of the *α* parameter, determines whether a slow (*p*_1_ = .01) or fast (*p*_2_ = 1) switcher wins (see Figure 4). This is because following a disaster, the initial growth of a genotype is determined by *R*(1 – *p*) – *α* which decreases with larger switching probabilities *p*. Indeed, for *p* = 1 this term is negative and the genotype drops in frequency for a short time. If disasters happen frequently and target the same phenotype then this *α* can be the dominant force in determining the winner.

**Figure 4:**
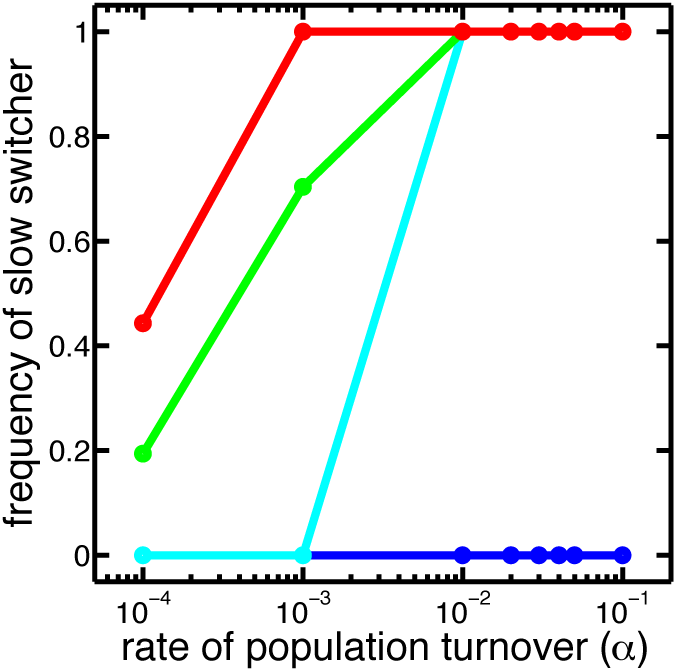
Population turnover determines the winner in a differential equation model. The frequency of the population composed of the slow switcher (*p* = .01) as opposed to the fast switcher (*p* = 1) is plotted against the rate of population turnover, *α*, in the differential equation model Eqn. 1. The different colors correspond to the number of disasters faced before the targeted phenotype is switched: 1 (blue), 5 (cyan), 10 (green), 25 (red). In this way the red line corresponds to a higher value of *t*_*c*_ than the blue. As the rate of population turnover increases, the slow switcher gains in frequency for all but the blue curve which corresponds to the lowest value of *t*_*c*_. The minimal value of population turnover that leads to the slow switcher winning decreases with longer durations in environments, i.e. higher *t*_*c*_ values.

We return to our stochastic model to weigh the competing pressures of competition and survival. In an environment of *t*_*c*_ = .99 a strategy of *p* = .01 always goes extinct when considered in isolation (Figure 1) and yet it is the best strategy for competition (Figure 2). The reason for this seeming contradiction is a significant difference in time scales (see Figure 5A). The time it takes *p* = .01 to go extinct is at least ten times greater than the time it takes to win a competition. Thus, while we expect *p* = .01 to go extinct eventually, it has enough time to outcompete faster switchers with *p* = 1.

**Figure 5:**
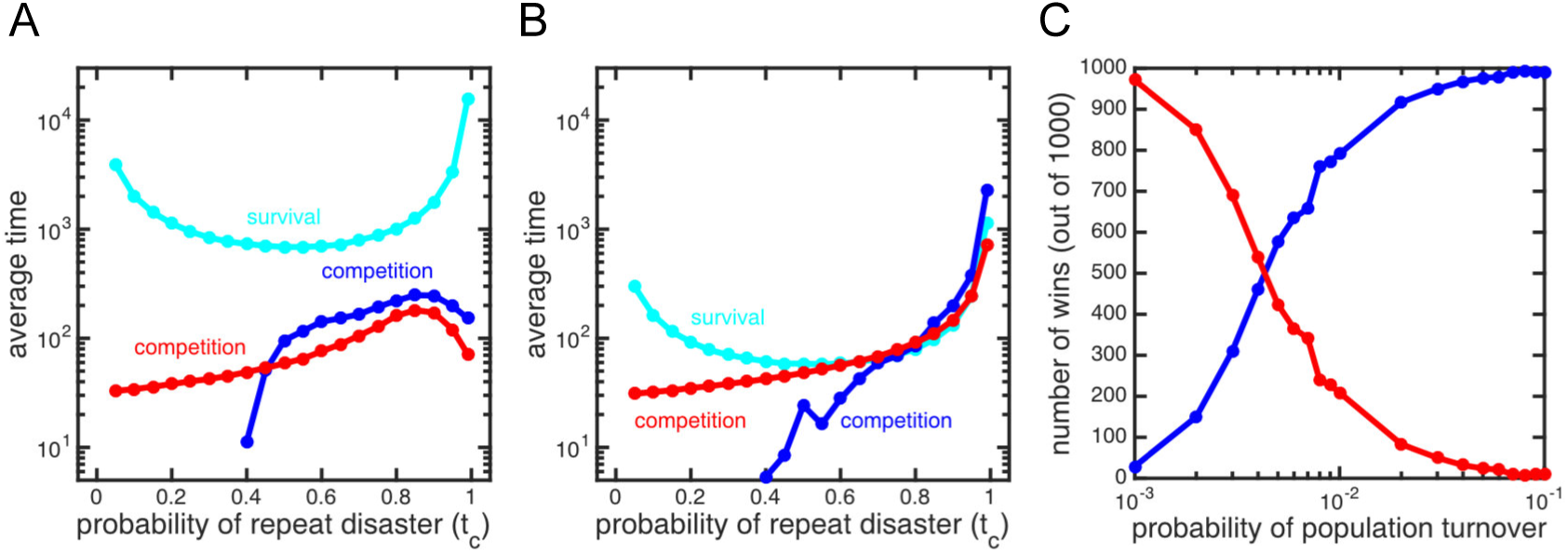
Time scale separation for survival and competition. **A**) The time scales over which the selective pressures of survival and competition act are shown as a function of the probability of repeat extinction. Each time is an average of 10000 simulations. The time for the slow switching strategy (*p* = .01) to go extinct (cyan) is at least ten times longer than it takes either *p* = .01 (blue) or *p* = 1 (red) to win in competition. The fast switching strategy did not go extinct and so is not plotted. **B**) The same as in A) except that the probability of population turnover is 100 times lower. The time scale for competition at *t*_*c*_ = .99 is now longer than the time scale for survival. This means that the *p* = .01 strategy will go extinct before winning the competition. **C**) The number of wins out of 1000 for *p* = .01 (blue) or *p* = 1 (red) switchers is shown as a function of population turnover when *t*_*c*_ = .99. The lower value of population turnover in B) is where the shorter survival times dominate and fast switchers win more often. As population turnover increases to the value in A), the survival time scale becomes longer allowing competition to dominate and the slower switcher to win more frequently.

The different time scales for competition and survival can be adjusted by changing parameters such as the rate of population turnover (see Figure 5B) or disaster probability (see Supplementary material). By decreasing the probability of population turnover (similar to *α* in the differential equation model), we can increase the time it takes for the slow switcher (*p* = .01) to win the competition against a fast switcher (*p* = 1). Similarly, the decreased value of population turnover gives less opportunities to diversify and so reduces the survival time. The net effect is that the survival time scale is shorter than the time scale for competition. As a result, the slow switcher goes extinct before winning the competition (see Figure 5C).

So far the cases considered have all been competitions between only two genotypes in an environment with fixed *t*_*c*_. To see if populations of potentially many genotypes can evolve to respond to the selective pressure imposed by the value of *t*_*c*_, we implement an evolutionary simulation in which genotypes mutate to give rise to new genotypes with a new characteristic switch rate. The mutation probability is 10^−3^ and a new switching probability is chosen from a uniform distribution from 10^−5^ to 1. We begin each simulation with a clonal population whose switch rate is *p* = .0001. The population goes through rounds of death and reproduction in a fixed environment (*t*_*c*_) for 100K iterations or until the population goes extinct. Figure 6A shows evolution within a *t*_*c*_ = .99 environment. The average probability of switching from 1000 simulations evolves to the optimal switching probability *p* = .01 but all go extinct prior to 80K rounds. This contrasts sharply with evolution in a *t*_*c*_ = .5 environment in which only 6 of 1000 simulations went extinct (Figure 6B). All other simulations lasted the entire duration with an average switching probability close to the optimal of *p* = 1.

**Figure 6:**
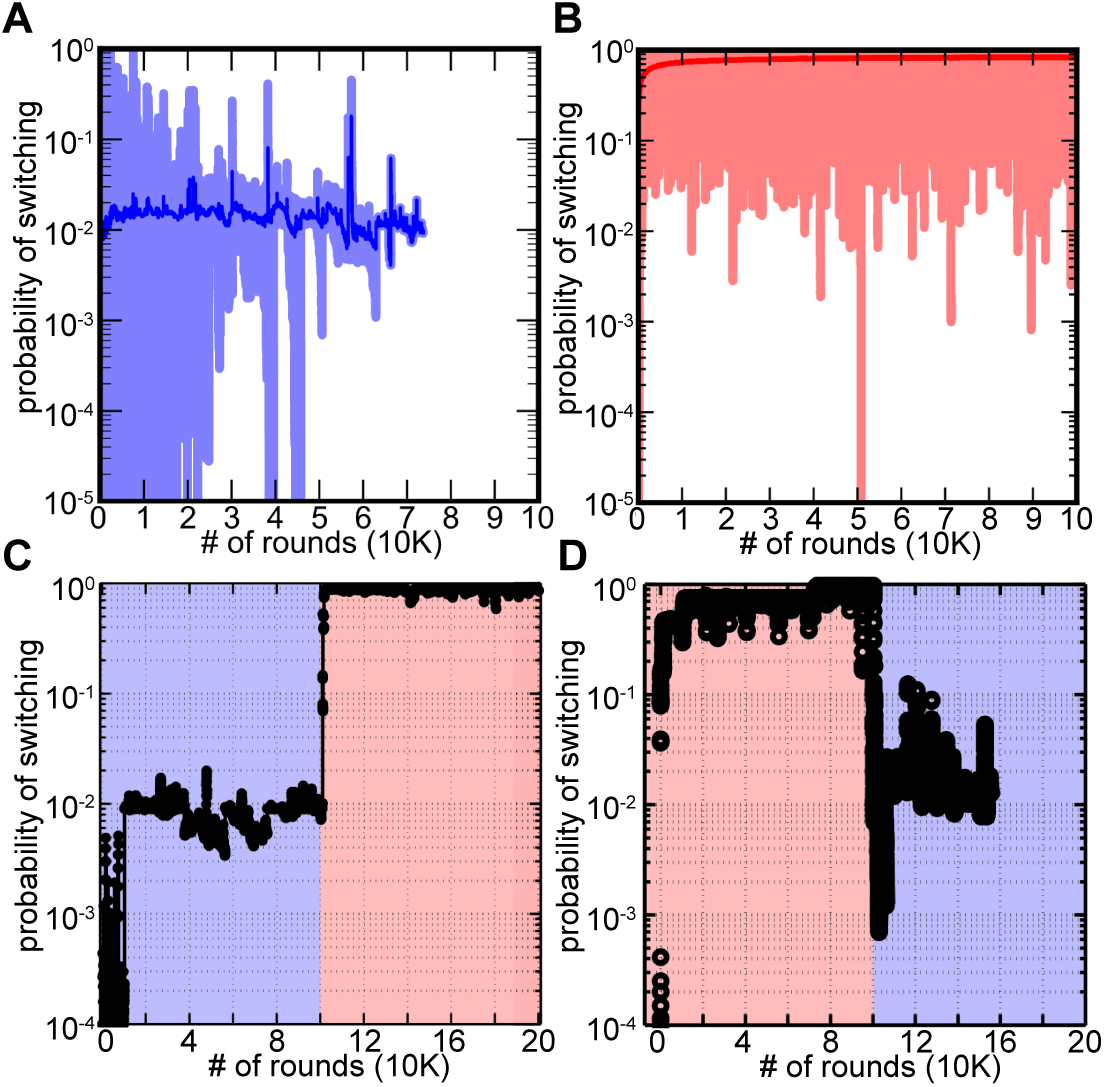
Evolution of switch rates in environments with different probabilities of repeat disasters. **A**) The average probability of switching over 1000 simulations of evolving populations (dark blue) is plotted over time in an environment with *t*_*c*_ = .99. The lighter area shows simulated populations removing the top and bottom 5%. Populations quickly evolve to *p* = .01 and then go extinct. **B**) Similar to A) but with *t*_*c*_ = .5 and a red scale for coloring. Populations evolve a probability of switching close to *p* = 1 and all but 6 out of 1000 survive the duration of the simulation. **C**) Similar to A) and B) but environments switch from *t*_*c*_ = .99 (blue) to *t*_*c*_ = .5 (red). It took many simulations to find a population that survived *t*_*c*_ = .99 but once it did, transfer to an environment with *t*_*c*_ = .5 saw the evolution of higher probabilities of switching close to the optimal *p* = 1 (average switch probability shown in black). **D**) Same as C) but in reverse order. The population average (black) evolves to the optimal competitive probabilities *p* = 1 in *t*_*c*_ = .5 (red) and *p* = .01 in *t*_*c*_ = .99 (blue) but ultimately goes extinct in *t*_*c*_ = .99.

The speed of adaptation and general results are further confirmed in Figures 6C and D when populations experience an environmental shift. Figure 6C shows the evolution of a population that survived *t*_*c*_ = .99 (a rare event) and is subsequently transferred to a different environment with *t*_*c*_ = .5. In *t*_*c*_ = .99, the population evolves to the optimal switch probability: *p* = .01. As the environment changes to *t*_*c*_ = .5, the population adapts by evolving to a switch rate close to the optimal of *p* = 1. The population remains at a high switch rate and survives for the rest of the simulation, 100K iterations. The reverse environmental fluctuation is shown in Figure 6D: a population evolving in *t*_*c*_ = .5 is transferred to an environment with *t*_*c*_ = .99. In this simulation, the population evolves to *p* > .7 in *t*_*c*_ = .5. When the environment shifts to *t*_*c*_ = .99, the population evolves to *p* = .01 and fluctuates before ultimately going extinct. In this scenario, the drive to respond to competition left the winning genotypes vulnerable to extinction from unpredictable environmental stress.

### Discussion

Stochastic phenotype switching is a canonical microbial bet-hedging strategy that increases fitness and long-term survival in unpredictable environments [6, 10, 13, 24, 34, 35]. However, the long time scales over which some bet-hedging strategies are manifest may conflict with other selective pressures. We uncover such an evolutionary tension between the timescales at which bet-hedging traits are adaptive. We study a mathematical model where environmental disasters select for diversification bet-hedging strategies, i.e. organisms switch between phenotypes to survive stochastic selection. We find that in environments where the type of disaster is positively correlated in time, switch rates that optimize competitiveness were favored over the short-term, while those that allow genotypes to avoid extinction were favored over the long-term. Since the time scale for competition is shorter than that for survival, populations evolve better competitive bet-hedging strategies, but this ultimately leaves them vulnerable to extinction.

The model presented in this paper has four key elements: genotypes that switch between phenotypes, disasters that annihilate a specific phenotype, population turnover caused by non-disaster death, and regrowth back to a carrying capacity. These elements are quite general and may appear in many real biological systems. Ecologically, one can think of our model as being a description of free-living bacteria that risk exposure to lytic phage capable of infecting only one of the two host phenotypes [45, 46], or bacteria living in a host risking detection by the immune system. These disasters occur stochastically, but the type of disaster (favoring either A or B cells) may be correlated in time. Furthermore, population expansion may be limited by the available space and resources found in the environment or host. Interestingly, in the bacteria-host scenario, if the host continually mounts a response against the most abundant bacterial phenotype then this would be similar to an environment *t*_*c*_ = 0 when “disasters”, i.e. immune responses, continually switch between targeted phenotypes. In this case, the survival probability of a switching organism actually increases when compared to environments with intermediate values of *t*_*c*_.

In the absence of temporal correlation of disasters, the strategy that maximizes short-term fitness also maximizes long-term survival (*p* = 1 in *t*_*c*_ = .5 wins in competitions and never goes extinct). In contrast, when risk is correlated in time, the strategy that maximizes short-term competitive fitness becomes one of the worst for long-term survival, and vice versa. Over short time periods selection favors traits that increase reproductive success. When risk is correlated in time slower rates of stochastic switching increase fecundity and competitive fitness because offspring of the opposite phenotype are likely to be killed by the next disaster. The downside of slower switching organisms is that they are far more susceptible to extinction (see Figure 1). Slower stochastic switching thus exhibits strong time-dependent fitness effects in our model, being advantageous over the short-term but costly over long time periods (see Figure 5). Importantly, in our simulations fast-switching strains capable of long-term survival in high *t*_*c*_ environments were driven extinct during competition by slower switching strains that then promptly went extinct following a disaster (see Figure 6).

Most prior models studying stochastic switching in fluctuating environments consider continuous populations that grow without limits [1, 8, 9, 32, 42]. This has two major effects relevant to the study of bet-hedging: first, it eliminates the risk of extinction caused by environmental fluctuations, and second, it decouples competing lineages so that the behavior of one lineage has no effect on the absolute fitness of competitors within the same population. We relax both of these constraints and find that imposing a carrying capacity on the population radically changes the ability for natural selection to favor the bet-hedging strategy that maximizes long-term fitness. By enforcing a lower limit on the probability of extinction, a carrying capacity allows extinction to play a powerful demographic role in shaping life history evolution. Without a carrying capacity, populations could expand to the point that extinction is no longer a threat.

In addition to affecting survival, carrying capacities are also instrumental in competition. By limiting opportunities for reproduction, a carrying capacity couples the fitness consequences of one strain’s switching strategy to its competitors. Specifically, the effects of employing one strategy determines the number of available reproductive events for the other strategy– either in the same round of growth, or a future round. As a result, strains with higher short-term fitness, but lower long-term fitness, can displace competitors (see Figure 6). These competitive effects should become more influential if one phenotype reproduces slower than another, as is the case with bacterial persistence [5]. Optimal fitness with bacteria persistence balances a trade-off between a fast-growing/antibiotic-susceptible phenotype and a dormant/antibiotic-resistant phe-notype [6]. Imposing a carrying capacity on a population of microbes hedging against antibiotic exposure via persistence would create an additional tradeoff in which production of dormant cells would limit the number of other cells that could be produced. These tradeoffs also exist in systems with regular, predictable environmental change as might be found in experimental populations of *Pseudomonas fluorescens* [43]. The only requirements are a limit to population growth and organisms that can produce more than one phenotype with different reproductive rates.

The stochastic phenotype switching considered in this paper fits within the hierarchical model of bet-hedging advanced by Andrew Simons [47]: stochastic switching itself is a primary bet-hedging trait that effectively improves fitness in unpredictable, fluctuating environments, while rapid switch rates can be a second-order bet-hedging trait beneficial only over long time periods. With this view, organisms that do not switch phenotypes are quickly driven extinct by environmental fluctuations. Long-term selection clearly favors switch rates rapid enough to avoid extinction, but this works against short-term selection for slow switching imposed by competition. Depending on the switch rates, there may be a significant time scale asymmetry between these selective agents. This asymmetry, coupled with the very real possibility of extinction during early-phase competition, may limit the ability of natural selection to favor higher-level bet-hedging strategies that maximize long-term fitness.

